# The prevalence of dementia in humans could be the result of a functional adaptation

**DOI:** 10.1101/2023.01.17.524364

**Authors:** Alan G Holt, Adrian M Davies

## Abstract

In this paper we propose that high copy number of the mitochondrial genome in neurons is a functional adaptation. We simulated the proliferation of deletion mutants of the human mitochondrial genome in a virtual mitochondrion and recorded the cell loss rates due to deletions overwhelming the wild-type.

Our results showed that cell loss increased with mtDNA copy number. Given that neuron loss equates to cognitive dysfunction, it would seem counter-intuitive that there would be a selective pressure for high copy number over low. However, for a low copy number, the onset of cognitive decline, while mild, started early in life. Whereas, for high copy number, it did not start until middle age but progressed rapidly. There could have been an advantage to high copy number in the brain if it delayed the onset of cognitive decline until after reproductive age. The prevalence of dementia in our aged population is a consequence of this functional adaptation.

## 1 Introduction

In a cell, copies of mitochondrial DNA (mtDNA) far outnumber those of nuclear DNA (nDNA). The mtDNA copy number varies between organs but it is particularly high in neurons where there can be several hundred to several thousand copies [11], compared to nDNA which has, usually, only two copies of each gene.

Previously we proposed that cells operate an energy feedback mechanism which enables cloning of mtDNA in the cell if there is an energy deficit and disables it if there is a surplus [18, 19]. We implemented this mechanism in a simulation model that we developed to study the proliferation of deletion mutations (mtDNA_*del*_) within neurons. While the energy feedback mechanism regulated the wild-type (mtDNA_*wild*_) it was possible for mtDNA_*del*_ to clonally expand, ultimately, resulting in cell death.

In this paper, we use our simulation model to study cognitive dysfunction due to neuron loss over a range of wild-type copy numbers (*CN*_*wild*_). We believe the results provide an insight into why *CN*_*wild*_ is high in neurons and propose that it is a functional adaptation. The results showed that cell loss increases with *CN*_*wild*_. Given that neuron loss is a significant factor in cognitive impairment [2], it would seem counter-intuitive that selection would favour high *CN*_*wild*_ over low. However, our results showed cognitive decline starts early in life, for low *CN*_*wild*_; anywhere from adolescence to early adulthood. Whereas for high *CN*_*wild*_, the onset of cognitive decline is delayed until middle age. Given that humans are now living much longer than we would have for most of our evolutionary history, dementia may be a byproduct of our early ancestors evolving a solution to cognitive decline in their reproductive years.

The contribution of this paper is three-fold:

- Creation of a simulated environment that models the proliferation of mtDNA_*del*_ within a virtual organelle and records the effects on the mtDNA_*wild*_ population.
- Studies the effect of *CN*_*wild*_ on cognitive decline due to the rate at which mtDNA_*del*_ proliferate and, ultimately, overrun mtDNA_*wild*_.
- Proposes that high *CN*_*wild*_ in neurons was an evolutionary advantage over low *CN*_*wild*_, despite the consequences of increasing the likelihood of dementia in later life.

## 2 Background

The most obvious differences between mtDNA and nDNA are their size and copy number. Human mtDNA contains 16,569 base pairs compared to nDNA with more than three billion. In a diploid cell, the nDNA consists of pairs of homologous chromosomes, thus, most genes will be present as two copies. In contrast, there are several hundred to several thousand copies of mitochondrial genes. While mtDNA copy number varies between organs, it is particularly high in neurons.

While the mtDNA and nDNA share a host cell they reside in distinct local environments. mtDNA are in close proximity to the electron transport chain and are exposed to free radicals generated as a byproduct of oxidative phosphorylation [7]. The nDNA is isolated from the electron transport chain and protected by the nuclear envelope.

mtDNA replicate independently of cell division [23]. The total mtDNA mass is regulated by the cell [27, 28], nevertheless, mtDNA constitute a population of replicons, potentially, capable of selfish behaviour [6]. While it is in the cell’s interest to maintain a population of mtDNA_*wild*_ in order to meet its energy requirements, continual population growth would be detrimental. Indeed, proliferation of mtDNA mutants is well documented and associated with neurodegenerative diseases [13, 12, 16, 30, 4].

We propose that mtDNA and nDNA have adopted different reproductive strategies akin to *r/K* selection [26] in response to the hazardous environment of the mitochondrial matrix [33]. Animal species that adopt an *r* strategy tend to be small, have a short lifespan and high reproductive rates. Whereas species that adopt a *K* strategy tend to be large, live long and reproduce slowly [32]. Thus mtDNA, with its high turnover rate resembles an *r*-like strategy while nDNA, which must match the lifespan the neuron, is *K*-like. Accordingly, nDNA is better protected and has superior repair pathways compared to mtDNA [37], while mtDNA mitigates damage to the population through attrition.

Mutation frequency increases with wild-type population size as the cell has to maintain a greater rate of cloning, resulting in an increased number of mutation events. That mtDNA copy number is high in neurons calls for explanation.

There are various theories for aging, the most well known is the mitochondrial free radical theory of aging [17], which proposes free radicals as causal agents in mutation. The single greatest risk factor for dementia is age [34]. Furthermore, the probable primary cause for the cognitive dysfunction observed in dementias, such as Alzheimer’s, is cell loss, where the greater the cell loss the more profound the cognitive dysfunction [2, 1]. It seems counter-intuitive that an adaptation would promote cognitive dysfunction.

A common reason given for high copy number is that it is dependent upon the energy demand of the tissue [31, 24, 9, 25, 3, 5]. Indeed, a cell has an energy requirement and the energy production is distributed amongst the mtDNA population. However, the graph in Fig. 1 shows that energy demand and copy number across different organ types is weakly correlated.

**Figure 1:**
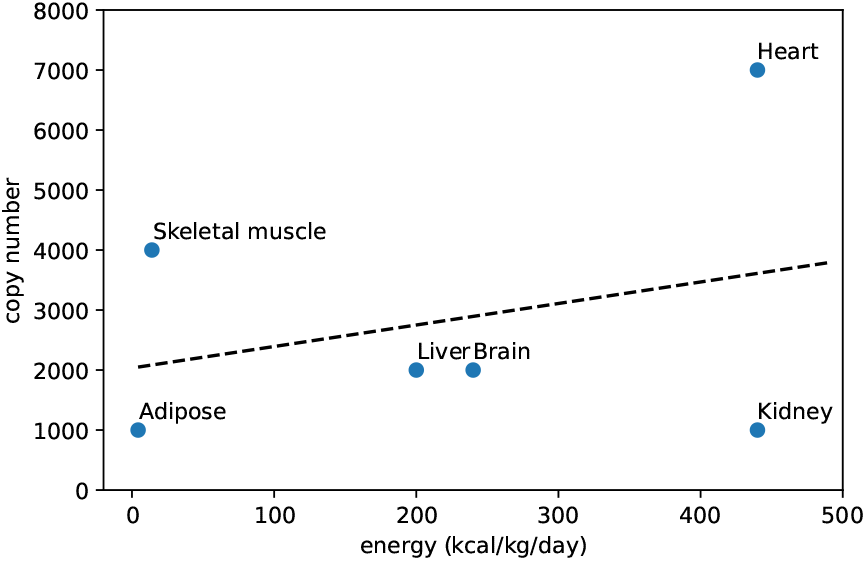
Per organ mtDNA copy number versus energy usage. Sources: [21, 38, 8, 20]

In some organs, like the kidney, the (per gram) high energy demand can be met with a small copy number of mtDNA_*wild*_, yet the brain, with a lower energy demand, requires a higher copy number. The reason for this is that copy number is likely dependent upon, not just the cell’s energy demand, but also the per mtDNA energy generation; as reflected in the equation below:

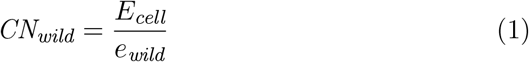

where *CN*_*wild*_ is the wild-type copy number, *E*_*cell*_ is a cell’s energy demand and *e*_*wild*_ is the per mtDNA_*wild*_ energy generation.

The per mtDNA_*wild*_ energy generation *e*_*wild*_ must, therefore, vary across organs. In which case, mtDNA_*wild*_ is capable of generating high *e*_*wild*_, thus meeting a cell’s energy demand with only a small copy number. Yet, neurons possess copy numbers ranging from several hundred [35] to several thousand [29].

## 3 Analysis

In this section we analyse the effect *CN*_*wild*_ has on neuron loss and the consequent cognitive dysfunction^1^. We developed a simulation model of mtDNA proliferation within a pseudo organelle based on the mitochondrion in neurons. We ran the simulation over a human lifespan (100 years) and noted cell loss rates. Cell loss rate is the ratio of cells that died before the host over the total cells:

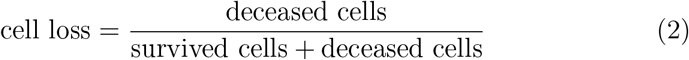

mtDNA proliferation within our simulator is governed by four basic principles, namely. *expiration replication mutation* and *competition*. We expand on the implementation of our simulation model^2^ below, but for a detailed description, see [19, 18].

mtDNA operate in a hostile environment and are subject to oxidative stress due to free radicals. Consequently, mtDNA *expire* with a half-life of 10-30 days [14, 22]. In our simulation we subject mtDNA to random *damage* pursuant to a 30 day half-life.

Given its short lifespan, mtDNA_*wild*_ must *replicate* by cloning in order to perpetuate the species. In our simulation, mtDNA clone at random according to a Bernoulli trial success of probability *P*_clone_ = 0.01 which is sufficient to counter the attrition rate and maintain a population. Otherwise, we assume replication is selfish and that it is up to the cell to control population level.

While it is in the cell’s interest to maintain a population of mtDNA, it must ensure the population does not grow out of control. The energy production in the cell is distributed amongst the mtDNA_*wild*_ population, see Eq (1). We implement an energy surplus/deficit feedback mechanism that suppresses cloning in the cell when there is an energy surplus and enables it when there is a deficit:

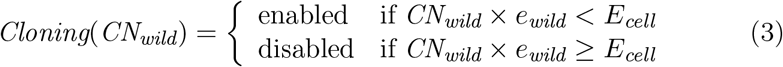

At random, we introduce errors into the cloning process which gives rise to mtDNA_*del*_. Like mtDNA_*wild*_, mtDNA_*del*_ replicate, however, mtDNA_*del*_ do not contribute to the cell’s energy requirements. While mtDNA_*del*_ are subject to cloning “on/off” signals from the cell, the cell’s energy feedback mechanism does not respond to mtDNA_*del*_ population levels (only mtDNA_*wild*_ population levels). Thus mtDNA_*del*_ can (and do) undergo clonal expansion.

We impose a capacity on the organelle which, if reached, prevents cloning of any mtDNA. Cloning can only resume after some attrition. When the capacity is reached, mtDNA enter into a *competition* for space. We call the moment at which the mtDNA population reaches this limit, the *competition point*.

Cloning is a sequential process, that is, an mtDNA cannot simultaneously clone multiple times and must complete its current cloning process before entering the next. The time to clone is proportional to an mtDNA’s size; given that mtDNA_*del*_ are smaller than mtDNA_*wild*_, deletions have a replicative advantage over wild-type.

If the mtDNA_*wild*_ loses this competition and is overwhelmed by mtDNA_*del*_, the cell dies. Cell death occurs when the mtDNA_*wild*_ population is not sufficient to meet the cell’s energy requirements. The decline in the mtDNA_*wild*_ population may take many years, but if the competition point is reached the mtDNA_*wild*_ rarely survives until the end of host’s lifetime.

For each *CN*_*wild*_ = (50, 100, 250, 500, 1000, 2000) we ran the simulation 100 times. We use resampling with replacement bootstrap methods [10] to generate a sampling distribution for the lifespan of the cell. Table 1 shows the parameters used in all the simulations. Only the *CN*_*wild*_ value is varied across simulation runs.

**Table 1:** Simulation parameters

| Parameter | Values |
| --- | --- |
| $P_{clone}$ | $0.01 h^{-1}$ |
| $P_{mutate}$ | $8 \times 10^{-4}/\text{clone event}$ |
| Capacity | 10,000 |
| Half-life | 30 days |
| mtDNA<sub>wild</sub> clone time | 2 $h$ |
| Simulation runtime | 100 years |

Previously, we used the cell loss threshold of 20% as the onset of dementia and 40% as advanced [19]. Below 20% was deemed healthy aging and, while the lifespan of the host was fixed at 100 years, it is unlikely any host survived beyond the point of 40% cell loss.

The graph in Fig. 2 shows cognitive decline for each value of *CN*_*wild*_. Cognitive decline occurs early in life for low copy number *CN*_*wild*_ *≤* 100, however, the decline is slow for the remaining lifespan. Cell loss does not exceed 20% until advanced old age. For high copy number, *CN*_*wild*_ *≥* 1000, cognitive decline is pushed back to middle age but dysfunction then progresses rapidly. The onset of dementia occurs in early old age and increases in severity in advanced old age.

**Figure 2:**
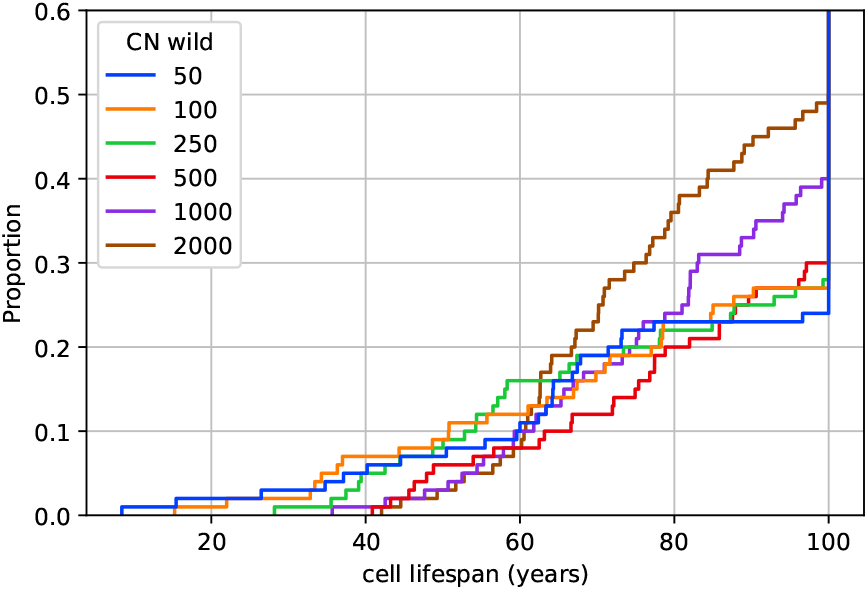
The proportion of cell loss over time for *CN*_*wild*_ = (50, 100, 250, 500, 1000, 2000). Cell loss equates to cognitive dysfunction. We deem 20% to be the onset of dementia and at 40%, dementia is advanced. Beyond 40% it is unlikely that the host survived despite the simulation running for the full 100 years.

The top graph in Fig. 3 shows the time at which deceased cells reached the competition point and the bottom graph shows how long after reaching the competition point, the cells survived. It can be seen that for lower copy number the competition point is reached much earlier compared to higher copy number. Furthermore, for low copy number, the cells are short lived once the competition point is reached.

**Figure 3:**
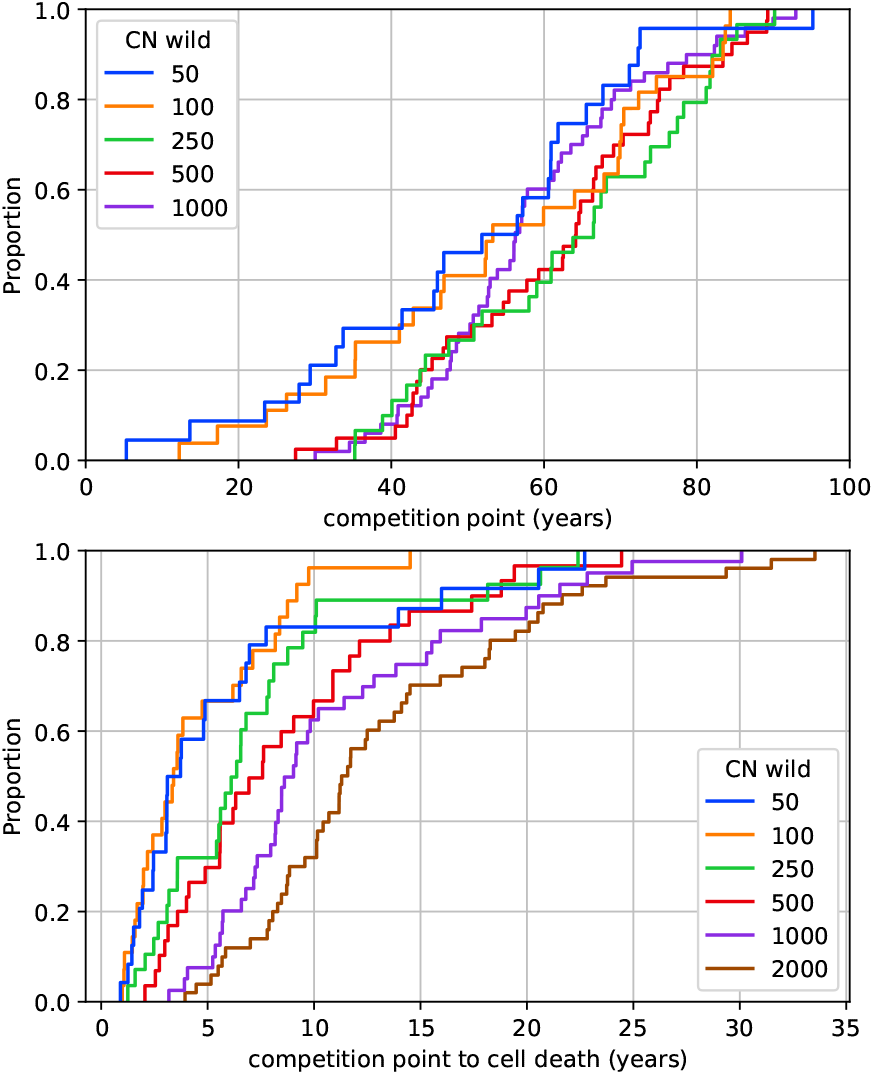
Top graph: proportion of deceased cells that reached the competition point at a given time. Bottom graph: how long deceased cells survived after the competition point had been reached.

Figure 4 shows the heteroplasmy for deceased cells. Heteroplasmy is much lower for low *CN*_*wild*_ meaning that there is more clonal expansion, that is, the mtDNA_*del*_ population is distributed amongst fewer species.

**Figure 4:**
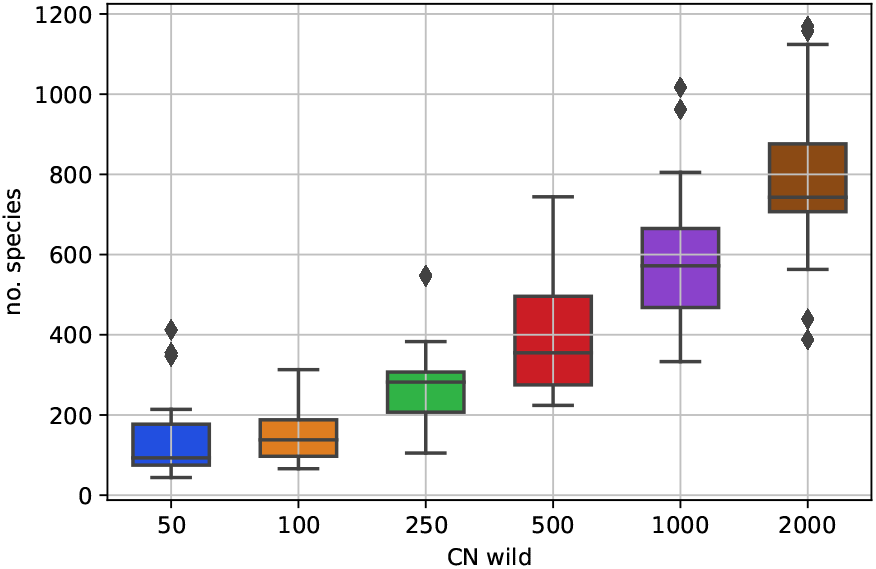
Heteroplasmy. The boxplot shows the number of mtDNA_*del*_ species throughout the life of the cell.

That cell loss increases with copy number is not a surprise as higher *CN*_*wild*_ (for a given mutation probability) would yield more mutation events. What was less expected was that cognitive decline occurs quite early in life for low *CN*_*wild*_. High copy number, while causing dementia in later life, appears to delay any cognitive decline until middle age. Thus, hosts that evolved a high mtDNA_*wild*_ copy number in neurons would have had an advantage during their reproductive years.

## 4 Discussion

The proximity of mtDNA to the electron transport chain means that it is subject to a higher rates of damage from free radical species than nDNA. As a population of short-lived replicons, mtDNA mitigates against damage through attrition. However, damage caused by free radicals also disrupt the cloning process which gives rise to species of mtDNA_*del*_ that compete with mtDNA_*wild*_. mtDNA_*wild*_ can be overrun by mtDNA_*del*_ resulting in cell loss and dysfunction. For mitotic cells, with a lifespan of weeks to months, mutant proliferation is not critical but for neurons, that need to last for many decades, some resilience to the expansion of mtDNA_*del*_ is essential.

That *CN*_*wild*_ is high in neurons may be considered an unfortunate consequence of the energy needs of the brain yet, the brain’s energy needs are moderate compared to some organs that have lower *CN*_*wild*_. This lack of correlation between copy number and energy demand by organ, as depicted in the graph of Fig. 1, merely tells us that linear regression is a bad model for predicting *CN*_*wild*_ as it involves only one variable, namely, cellular energy demand.

We model *CN*_*wild*_ as dependent upon two variables, that is, cellular energy demand *E*_*cell*_ and per mtDNA_*wild*_ energy production *e*_*wild*_ (Eq (1)). Indeed, the graph in Fig. 1 implies that *e*_*wild*_ varies across organs and that, in each organ, energy utility of mtDNA_*wild*_ was able to evolve independently.

We suggest that elevation of *CN*_*wild*_ in the brain was in response to the neural mtDNA_*wild*_ lowering its energy output. An increase in the rate of oxidative phosphorylation could have been achieved by greater transcription and translation of the mtDNA encoded genes, thus, obviating the need to raise *CN*_*wild*_. However, as *CN*_*wild*_ *is* high in neurons, it suggests that this is not what happened. Rather the cell balanced its energy demand with the reduction in mtDNA_*wild*_ energy production by increasing *CN*_*wild*_. Furthermore, this was a functional adaptation that yielded a selective advantage to the gene that an increase to the rate of oxidative phosphorylation lacked.

There is a difference in mtDNA proliferation between low and high copy number. The early cognitive decline caused by low *CN*_*wild*_, while small, may have been enough to have been an evolutionary disadvantage and, therefore, selected against in favour of high *CN*_*wild*_.

However, there is a consequence for high *CN*_*wild*_. While the start of cognitive decline is pushed back to middle age, it does so at the expense of dementia in later life. Furthermore, the severity of that dementia correlates with the increase in *CN*_*wild*_. Thus, a host’s predisposition to dementia could be linked to wild-type copy number.

Functional adaptations occur for the benefit of the gene not necessarily the host, thus, consequences of high *CN*_*wild*_ would likely escape the attention of evolution given the onset of dementia occurs after reproductive age.

## 5 Conclusions

In this paper we simulated the proliferation of mtDNA_*del*_ in neurons over a range of mtDNA_*wild*_ copy number 50 ≤ *CN*_*wild*_ ≤ 2000. We recorded the cell loss rates, as loss of neurons is a primary cause of dementia [2, 1]. The results showed that cell loss increased with *CN*_*wild*_.

For low copy number *CN*_*wild*_ ≤ 100, cognitive decline begins in adolescence and rises to a small but appreciable level during early adulthood. Loss rates that would equate to dementia do not occur until advanced old age (*>* 80 years). In actual patients there is marked increase in the incidence of dementia after 65 with mortality at 7-10 years after diagnosis [36].

Dementia is a disease of old age but it is far more prevalent than is predicted at low *CN*_*wild*_ by our simulation. Furthermore, at low *CN*_*wild*_, our simulation predicts a gradual decline which does not match clinical observations. Rather, the onset and progression of dementia better matches our results for high copy number,namely, cognitive decline ranging from healthy aging to chronic dementia beginning after reproductive age. These results align with current observations that mtDNA copy number are in the many hundreds to low thousands. Furthermore, it could explain why some people are more predisposed to dementia than others given the range of observed copy number.

Given that genes behave selfishly, it is conceivable that *CN*_*wild*_ in neurons could have increased over evolutionary time. By lowering *e*_*wild*_ (Eq (1)) over generations, mtDNA_*wild*_ exploited the energy surplus/deficit feedback mechanism. Thus ensuring that the neurons maintain a high copy number in order to meet its energy requirements, despite the increase in the risk of dementia in later life,

Classically, the force of selection declines with age [15], and dementia is the consequence of the selective advantage of delaying cognitive decline until post-reproductive age. Therefore, we propose that the prevalence of dementia in old age is the result a function adaptation that increased the copy number of wild-type mtDNA in neurons.

## Footnotes

1 The data for this paper can be found at: https://www.kaggle.com/alanholt/copynumber-analysis.

2 The source code for the simulation is available in the Github repository: git@github.com:agholt/closs.git.

